# Genetic diversity of *Anopheles coustani* in high malaria transmission foci in southern and central Africa

**DOI:** 10.1101/2020.04.04.020537

**Authors:** Ilinca I. Ciubotariu, Christine M. Jones, Tamaki Kobayashi, Thierry Bobanga, Mbanga Muleba, Julia C. Pringle, Jennifer C. Stevenson, Giovanna Carpi, Douglas E. Norris, for the Southern and Central Africa International Centers of Excellence for Malaria Research

## Abstract

Despite ongoing malaria control efforts implemented throughout sub-Saharan Africa, malaria remains an enormous public health concern. Current interventions such as indoor residual spraying with insecticides and use of insecticide-treated bed nets are aimed at targeting the key malaria vectors that are primarily endophagic and endophilic. While these control measures have resulted in a substantial decline in malaria cases and continue to impact indoor transmission, the importance of alternative vectors for malaria transmission has been largely neglected. *Anopheles coustani,* an understudied vector of malaria, is a species previously thought to exhibit mostly zoophilic behavior. However, recent studies from across Africa bring to light the contribution of this and ecologically similar anopheline species to human malaria transmission. Like many of these understudied species, *An. coustani* has greater anthropophilic tendencies than previously appreciated, is often both endophagic and exophagic, and carries *Plasmodium falciparum* sporozoites. These recent developments highlight the need for more studies throughout the geographic range of this species and the potential need to control this vector. The aim of this study was to explore the genetic variation of *An. coustani* mosquitoes and the potential of this *Anopheles* species to contribute to malaria parasite transmission in high transmission settings in Nchelenge District, Zambia, and the Kashobwe and Kilwa Health Zones in Haut-Katanga Province, the Democratic Republic of the Congo (DRC). Morphologically identified *An. coustani* specimens that were trapped outdoors in these study sites were analyzed by PCR and sequencing for species identification and blood meal sources, and malaria parasite infection was determined by ELISA and qPCR. Fifty specimens were confirmed to be *An. coustani* by the analysis of mitochondrial DNA cytochrome *c* oxidase subunit I (COI) and ribosomal internal transcribed spacer region 2 (ITS2). Further, maximum likelihood phylogenetic analysis of COI and ITS2 sequences revealed two distinct phylogenetic groups within this relatively small regional collection. Our findings indicate that both *An. coustani* groups have anthropophilic and exophagic habits and come into frequent contact with *P. falciparum,* suggesting that this potential alternative malaria vector might elude current vector controls in Northern Zambia and Southern DRC. This study sets the groundwork for more thorough investigations of bionomic characteristics and genetic diversity of *An. coustani* and its contribution to malaria transmission in these regions.

## Introduction

Malaria is transmitted to humans by the infectious bite of female mosquitoes of the *Anopheles* genus, and approximately 70 *Anopheles* species are potential vectors of malaria that transmit the disease to humans effectively worldwide (Sinka et al. 2012). Species are often characterized as primary or secondary vectors: primary vectors are those mosquitoes that are abundant, most commonly feed on humans, and have measurable sporozoite rates, while secondary or alternative vectors can be uncommon, have low sporozoite rates, but may still play a role in malaria transmission (Oaks SC Jr. 1991).

Zambia, in sub-Saharan Africa, is a malaria-endemic country that has experienced high mortality and morbidity from this disease for decades (Mukonka et al. 2014, Ministry of Health Zambia 2015). Significant strides have been made to reduce malaria transmission, largely due to the implementation of vector control interventions (Bhatt et al. 2015). These interventions include vector control through the distribution of long-lasting insecticide-treated nets (LLINs), indoor residual spraying (IRS), treatment through intermittent preventive treatment in pregnancy (IPTp), and case management through the use of rapid diagnostic tests (RDTs) and artemisinin-combination therapy (ACT) (Chizema-Kawesha et al. 2010, Sutcliffe et al. 2012, MIS 2019, PMI 2019a). Despite successful reductions in morbidity and mortality, malaria remains endemic with over 6 million reported cases in 2018 (MIS 2019, WHO 2019a). While the scaling-up of malaria interventions such as widespread coverage by LLINS and IRS reduced transmission and parasitemia throughout many parts of Zambia, the disease continues to be a significant public health concern, especially in the northern region where Nchelenge District, Luapula Province, is recognized as a high transmission focus (Chanda et al. 2013, Mukonka et al. 2014, Nambozi et al. 2014, Hast et al. 2019). This region of Zambia reports over 350 confirmed cases per 1000 population (Moss et al. 2012, PMI 2019a, WHO 2019a). This has raised doubts about whether the progress made across Zambia could be maintained and called for more enhanced and targeted interventions, especially in northern Zambia (Kamuliwo et al. 2013).

Like Zambia, the Democratic Republic of the Congo (DRC) is a malaria-endemic country in the central region of sub-Saharan Africa, located northeast of Zambia, and in which malaria is a leading cause of mortality and morbidity, accounting for approximately 12% of malaria cases and 11% of deaths worldwide (Messina et al. 2011, Stone et al. 2015, WHO 2019a). Despite actions taken to scale-up interventions in the DRC, such as the distribution of LLINs with over 50% household coverage throughout the country, malaria transmission still remains high and progress appeared to have stalled according to the world malaria health reports of 2017 and 2018 (Koukouikila-Koussounda and Ntoumi 2016, WHO 2019a). This country was part of the launch of the WHO and RBM Partnership to End Malaria in 2018 through a high burden to high impact country-led approach in the hopes of continuing progress and reaching the 2025 goals of the Global technical strategy for malaria (WHO 2015, 2019b). In 2018, the DRC was one of six countries that accounted for more than half of all malaria cases, globally, and had an estimated 26 million cases of malaria (WHO 2019a). Moreover, in this country, malaria remains the leading cause of morbidity and mortality, and accounts for 19% of deaths among children under the age of 5 (Ferrari et al. 2016, PMI 2019b).

Zambia and the DRC exhibit seasonal transmission that follows rainfall patterns, in which malaria peaks after the rains when mosquito populations increase (Masaninga et al. 2013). However, in Nchelenge District, malaria transmission is intense with limited seasonal fluctuations (Mharakurwa et al. 2012). In Nchelenge, the primary vectors of malaria in both the dry and wet seasons have been found to be *An. funestus* s.s. and *Anopheles gambiae* s.s. (Das et al. 2016, Jones et al. 2018, Hast et al. 2019). Malaria is also holoendemic in the DRC which borders Zambia to the north, but much less is known about malaria vectors and their phenology, other than that *An. funestus* s.s., *An. gambiae* s.s., and *An. coluzzii* are major vector species throughout the region, with the first two species exhibiting high biting rates (Bobanga et al. 2016, Nardini et al. 2017, Wat’senga et al. 2018). Vector control methods such as IRS and LLINs, which have been implemented throughout all of Zambia, and to a much less extent in the DRC, are aimed at targeting these vectors preferentially. In addition to a suboptimal coverage of vector control in northern Zambia and southern DRC, malaria may remain intractable due to the presence of alternative vector species that have largely remained unrecognized.

The focus on control and elimination methods for the well-recognized endophagic vector species highlights the fact that alternative vectors are rarely considered in existing malaria control programs, and are thought of as negligible because of their often zoophilic behavior (Fornadel et al. 2011). However, it has been observed that after primary vectors are reduced in a population, alternative vectors have the potential to sustain malaria transmission (Antonio-Nkondjio et al. 2006). Previous studies have indicated the presence of *P. falciparum* parasites in these alternative vectors in Kenya, Ethiopia, Zambia, and other regions in Africa (Stevenson et al. 2012, Degefa et al. 2015, Lobo et al. 2015, Nepomichene et al. 2015a, St. Laurent et al. 2016, Stevenson et al. 2016a). One of these alternative vectors, *An. coustani,* is a species previously reported to exhibit mostly zoophilic behavior (Gillies and DeMeillon 1968). However, recent studies from countries in southern Africa are bringing to light the potential contribution of this species to malaria transmission. In Zambia, this species displays an unexpectedly high degree of anthropophilic tendencies (Fornadel et al. 2011). In Kenya, this vector is both endophagic and exophagic and is thought to play a major role in outdoor malaria transmission (Mwangangi et al. 2013). In Madagascar, this vector has been shown to carry *P. falciparum* infections in both indoors and outdoors collections, and more recently, to act as a major local vector even though it is was previously a suspected alternative vector (Nepomichene et al. 2015b, Goupeyou-Youmsi et al. 2019). These findings emphasize that mosquito species such as *An. coustani* may contribute more significantly to malaria transmission than previously recognized. These findings warrant further study of these alternative vectors, including foraging behaviors, ecology, genetics, and potential roles in the transmission of malaria.

As anophelines commonly exist as species complexes, integrating molecular and phylogenetic analysis to collections of *An. coustani* enable the confirmation of species identification, and allow for further assessment of genetic diversity and relatedness within and between species complexes. This study focused on assessing the potential role *An. coustani* may play in *P. falciparum* transmission in northern Zambia and southern DRC by analyzing phylogenetic relationships and human exposure. Improved species identification and correct association of species with phenotypes relevant to vectorial capacity will allow for the design of better control strategies.

## Methods

### Mosquito collection and handling

As part of the International Centers of Excellence for Malaria Research (ICEMR) in Southern and Central Africa, mosquito specimens were collected solely outdoors from Nchelenge District, northern Zambia, using standard Centers for Disease Control and Prevention (CDC) light traps, and from two villages (Kilwa and Kashobwe) in Haut-Katanga Province, southern DRC, using CDC light traps and pyrethrum spray catches (PSC) (Figure 1). In Nchelenge District, Zambia, CDC light traps were placed overnight in the three following scenarios: outdoors where humans congregate, outdoors next to animal pens, and outdoors with a commercial human analogue bait (BG Lure^®^, BioGents). Households were selected using a similar sampling frame to that of other studies under the ICEMR program (Pinchoff et al. 2015, Stevenson et al. 2016b). Collections were performed over a 2-week period in August 2016 at eight households (4 inland along a stream and more than 3 km from Lake Mweru and 4 lakeside, close to Lake Mweru) for a total of 74 trap nights. Using a Latin Square design, trap scenarios were rotated through each household, such that each treatment occurred in each household at least once. Traps were activated at 6pm and tied shut and retrieved the following morning.

**Figure 1.**
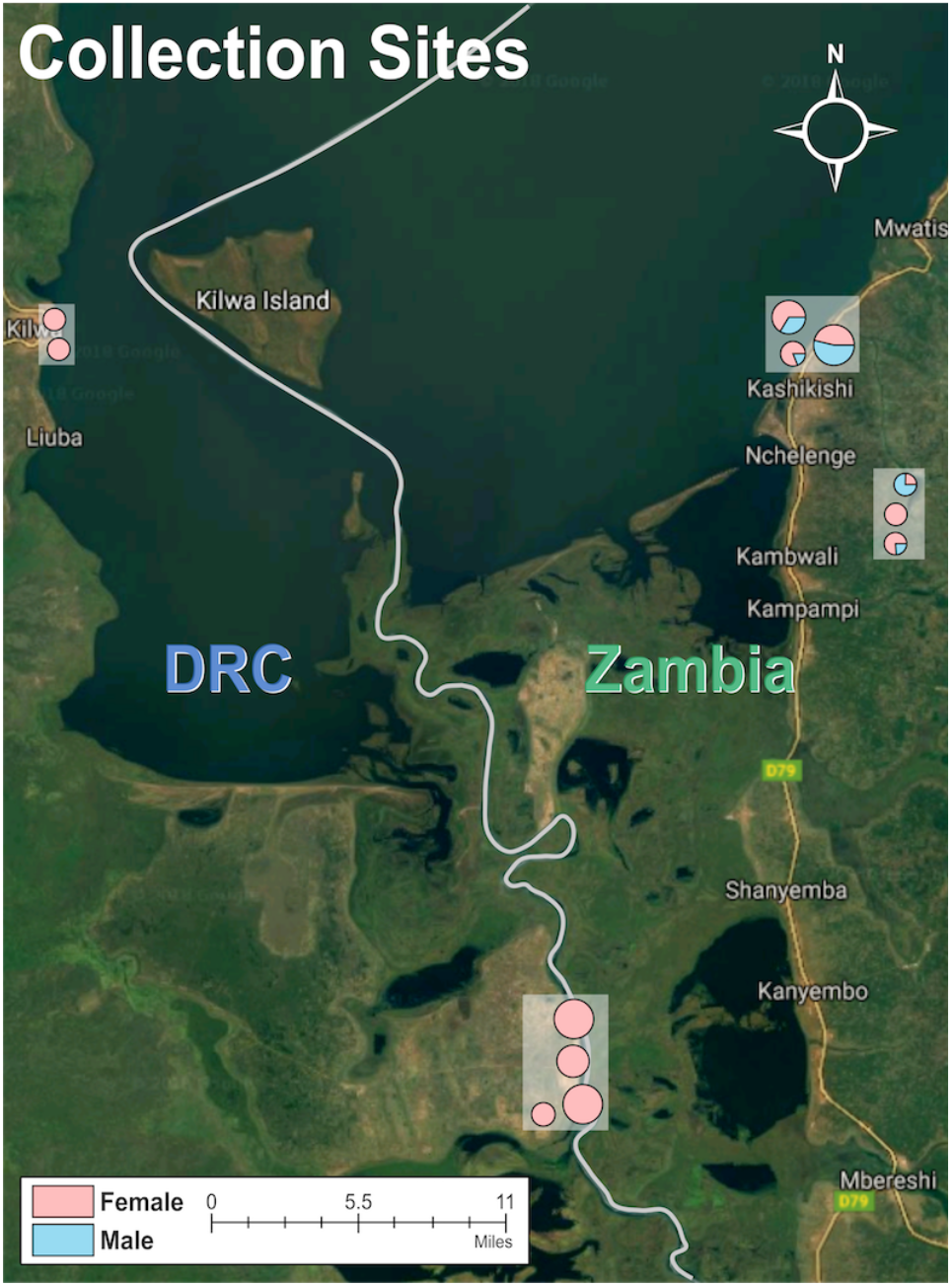
Distribution of caught mosquitoes over multiple sites in Zambia and the DRC. The size of the circle refers to the proportion of samples caught in a specific area relative to the size of the collection, and the color represents the ratio of males/females in a specific area.

In the DRC, the Kilwa Health Zone is located near Lake Mweru, across the lake from Nchelenge, Zambia, and the Kashobwe Health Zone is located near the Luapula River as it leaves the south end of Lake Mweru, providing abundant vector breeding sites throughout the year. In the Kilwa and Kashobwe Health Zones, 60 study households (Kilwa = 30 households, Kashobwe = 30 households) were randomly selected from village household census. Mosquito collections were performed by one of three collection scenarios: hanging CDC light traps indoors overnight next to a home occupant sleeping under a LLIN, hanging CDC light traps outdoors overnight by a window, or PSC early in the morning.

Anophelines collected in Nchelenge were killed by freezing, while those from the DRC were left at room temperature before being packaged. Anophelines were identified by sex and morphology with the aid of a dissecting microscope and dichotomous key at all field sites (Gillies and Coetzee 1987). Mosquitoes were then each placed individually in 0.6 mL microcentrifuge tubes that contained silica gel desiccant and a cotton wool plug. They were transported and stored at room temperature until processed in the laboratory at Johns Hopkins Bloomberg School of Public Health in Baltimore, Maryland.

### Isolation of DNA and Molecular Processing

In the laboratory, the abdomen of each collected anopheline was separated from the head and thorax using sterile forceps, and then stored in separate tubes at −20°C. Genomic DNA was extracted from the frozen mosquito abdomens with a salt extraction method as previously described (Post et al. 1993, Norris et al. 2001, Das et al. 2016). A fragment of the mitochondrial cytochrome *c* oxidase subunit I (COI) gene used for the Barcode of Life Database (BOLD), a molecular target that has been previously used for phylogeny construction of anophelines, was amplified and sequenced from specimens that were morphologically identified as *An*. *coustani* for a total of 50 specimens (Beebe 2018). The 698 bp BOLD fragment of the COI gene was amplified using LCO1490 and HCO2198 primers as previously published (Lobo et al. 2015). The 25 μl PCR mixture consisted of 2.5 μ1 of 10x buffer, 2.5 mM dNTP mixture, 30 pmol each of the forward and reverse primers, 2.0 U of *Taq* DNA polymerase (Invitrogen, Carlsbad, CA), and 1 μl of mosquito DNA template. The thermocycler (MultiGene™ OptiMax Thermal Cycler, Labnet International, Inc., Edison, NJ) conditions were identical to the ones described by Lobo and colleagues (2015).

A 750 bp fragment of the ribosomal DNA internal transcribed spacer region 2 (ITS2), a nuclear gene located between the 5.8S and 28S large subunit RNA genes that is often used for species verification and identification in anophelines, was also targeted for sequencing (Mohanty et al. 2009, Norris and Norris 2015). The ITS2 region was amplified from genomic DNA using ITS2A and ITS2B primers, and the 25 μl PCR mixture was identical to that for the BOLD fragment amplification PCR (Lobo et al. 2015). The thermocycler conditions were identical to the ones described by Lobo and colleagues (2015).

The PCR-amplified products of all specimens were visualized by electrophoresis on a 2% agarose gel. Resulting PCR products were then purified using the QIAquick^®^ PCR Purification Kit (Qiagen, Hilden, Germany) before Sanger sequencing by the Sequencing Facility at the Johns Hopkins School of Medicine. A multiplexed PCR was performed on DNA extracted from mosquito abdomens to detect multiple blood meals by a protocol that differentiates between possible mammalian host blood in female mosquitoes as animal-specific products (human, cow, dog, pig, goat) amplified from the cytochrome *b* mitochondrial gene as described in detail elsewhere (Kent and Norris 2005).

### Species assignment and Phylogenetic Analyses

For both COI and ITS2 targets, forward and reverse sequences for each sample were trimmed to remove ends with low Phred quality and then high-quality trimmed forward and reverse sequences were aligned to generate a single consensus sequence for each individual sample using Geneious (Biomatters, Auckland, New Zealand) version 11.1.5 (https://www.geneious.com). Each COI and ITS2 consensus sequences was then queried against the NCBI database using BLASTn for molecular species identification/assessment (Altschul et al. 1997). Samples were confirmed as a particular species when the COI BLAST results indicated that there was a minimum nucleotide identity of greater than 90% and a significant E-value <1×10^-5^. Full consensus sequences for each sample were submitted to GenBank and assigned accession numbers (Supplementary Table 1).

COI sequences that were confirmed as *An. coustani* after comparison with NCBI BLASTn were combined to generate a multiple sequence alignment using the MUSCLE algorithm and default parameters in the Geneious version 11.1.5 aligner. The multiple sequence alignment of 50 *An. coustani* sequences was then trimmed to a final length of 552 bp for COI and 662 bp for ITS2. Furthermore, a multiple alignment was then created with the confirmed *An. coustani* samples and other *Anopheles* species from the Hyrcanus and Coustani groups from the NCBI database for the COI target, and with the confirmed *An. coustani* samples and other *An. coustani* ITS2 sequences for the ITS2 target, as this latter gene region is not highly conserved between species.

Phylogenetic analysis was conducted using maximum likelihood (ML) inference as implemented in MEGA X (Kumar et al. 2018, Stecher et al. 2020). The evolutionary history was inferred by the Maximum Likelihood method and General Time Reversible model (Nei and Kumar 2000). One tree with the highest log likelihood for each target was included in this manuscript. Brach support was included by bootstrap with 1000 replications. The same steps were performed for analysis of the ITS2 sequences from the 50 *An. coustani* mosquitoes.

### Detection of *Plasmodium falciparum*

Head and thoraces of female *An. coustani* mosquitoes were homogenized in a phosphate-buffered saline based solution in 1.5 mL microcentrifuge tubes for enzyme-linked immunosorbent assay (ELISA) and genomic DNA was extracted from half of the homogenate using the Qiagen: DNeasy Blood and Tissue Kit protocol (Qiagen, Hilden, Germany) (Beier 2002, Gomes et al. 2017). As quality control, genomic DNA concentration of each extract was quantified using High Sensitivity double-stranded DNA (dsDNA HS) assay on a Qubit 2.0 Fluorometer (Life Technologies, Grand Island, NY). All female *An. coustani* head and thoraces were subjected to the ELISA assay which uses monoclonal antibodies (CDC, Atlanta, USA) targeting the circumsporozoite protein (CSP) of *P. falciparum* sporozoites (Burkot et al. 1984). In addition, the genomic DNA extracts from the same individual mosquitoes were screened for *P. falciparum* DNA using a SYBR Green qPCR assay that targets an 85 bp fragment of the *P. falciparum* lactate dehydrogenase *(Pfldh)* gene using primers described elsewhere (Parr et al. 2016). The 25 μl PCR mixture consisted of 12.5 μl SYBR Green PCR Master Mix (Life Technologies, Warrington, United Kingdom), 1 μM each of the forward and reverse primers, and 4 μl of template DNA. Each reaction was performed in a StepOne Real-Time PCR System (Applied Biosystems, Foster City, CA) and the cycling conditions were the following: 50°C for 2 minutes, 95°C for 10 minutes, followed by 45 cycles of denaturation at 95°C for 15 seconds, and annealing at 60°C for 1 minute. All samples were replicated in each reaction plate, no template controls (NTCs) were included and run alongside standard dilutions of gDNA of *P. falciparum* NF54 strain (1-10^5^ *P. falciparum* genome equivalent/μl).

### Genetic Diversity

DNAsp (version 5.10.1) was used to assess diversity and polymorphism statistics on the sequences obtained from the mosquito samples collected in Zambia and the DRC (Librado and Rozas 2009). The number of polymorphic sites (S), the number of haplotypes, average nucleotide diversity (nucleotide differences per site based on pairwise comparisons among DNA sequences) (*π*), mean number of nucleotide differences (k), genetic diversity (*θ*), and haplotype diversity (H_d_) (probability that two samples randomly sampled are unique) were all calculated for both COI and ITS2 sequences (Nei 1987).

## Results

A total of 42 female (31 from northern Zambia and 11 from southern DRC) and 15 male (northern Zambia) morphological *An. coustani* were caught outdoors as part of this study, representing less than 5% of all mosquitoes captured from these study sites as part of a larger collection. Morphological identifications were conducted on all 57 of these specimens at the field sites, and all species included in this study were identified as *An. coustani*. Following extractions in the laboratory, both a 698 bp (BOLD) COI fragment and a 750 bp ITS2 fragment were amplified from the abdomen of all samples. 100% of the males successfully amplified with these PCRs, while 7 (16.7%) of the female mosquito amplifications failed after multiple attempts; thus, 50 samples were included in the remaining assays and analyses. Following Sanger sequencing and comparison to NCBI databases, all 50 of these mosquitoes that were morphologically identified as *An. coustani* were molecularly confirmed. All 50 *An. coustani* COI sequences matched (with at least 90% nucleotide identity and a significant E-value <1×10^-5^) to previously reported *An. coustani* sequences deposited in NCBI (as of January 18, 2020). For the ITS2 fragment, 29 (58%) of these samples did not yield any close matches (closest match of ~80% was *Anopheles yatsushiroensis,* a member of the Hyrcanus Group that is widely distributed in Oriental and Palaearctic areas) when their consensus sequences were blasted on the NCBI databases, which is presumably due to the relative lack of data for this locus and species in the database (Fang et al. 2017). There are over twenty *An. coustani* COI sequences publicly available (NCBI BLASTn, January 2020) and this gene is highly conserved, with little interspecies variation (Margoliash 1963). In contrast, there are currently only three previously submitted sequences in NCBI GenBank for the less conserved ITS2 gene for *An. coustani* (Coleman 2007).

Of the 35 female *An. coustani* samples, 4 (10.8%) had blood meals identified solely as goat, 3 (8.6%) had blood meals identified solely as human, and 3 (8.6%) had mixed blood meals identified as goat and human. None of the 35 female anopheline samples were CSP ELISA positive. All 35 female *An. coustani* samples were also analyzed by qPCR for *Pfldh.* Six samples (17.1%, 95% CI 6.5-33.6%) yielded positive signals for *P. falciparum* (parasite load was in the range of 1 parasite/μl – 5 parasite/μl), of which 2 were blooded with human blood.

The Maximum Likelihood analysis was performed for both the COI and ITS2 fragments. The resulting phylogenetic tree that was constructed for the COI fragment (Figure 2) revealed that all specimens that were morphologically identified as *An. coustani* clustered together. The COI clustering revealed two well-supported molecular groups (hereafter “*An. coustani* group A” and “*An. coustani* group B”) with 74% and 99% bootstrap support, respectively, among the *An. coustani* samples. *An. coustani* group A encompasses 21 samples from the field and previously described *An. coustani* sequences from Zambia (KR014841 and KR014843), Mali (MK585958), and the Republic of Guinea-Bissau (KM097027). Within group A, there is substructure that does not reflect geography, with samples from both Zambia and the DRC grouping closely with the aforementioned samples published from other parts of Africa. *An. coustani* group B, which has 100% bootstrap support, contains the remaining 29 newly reported sequences from this collection, and has two well-supported groups.

**Figure 2.**
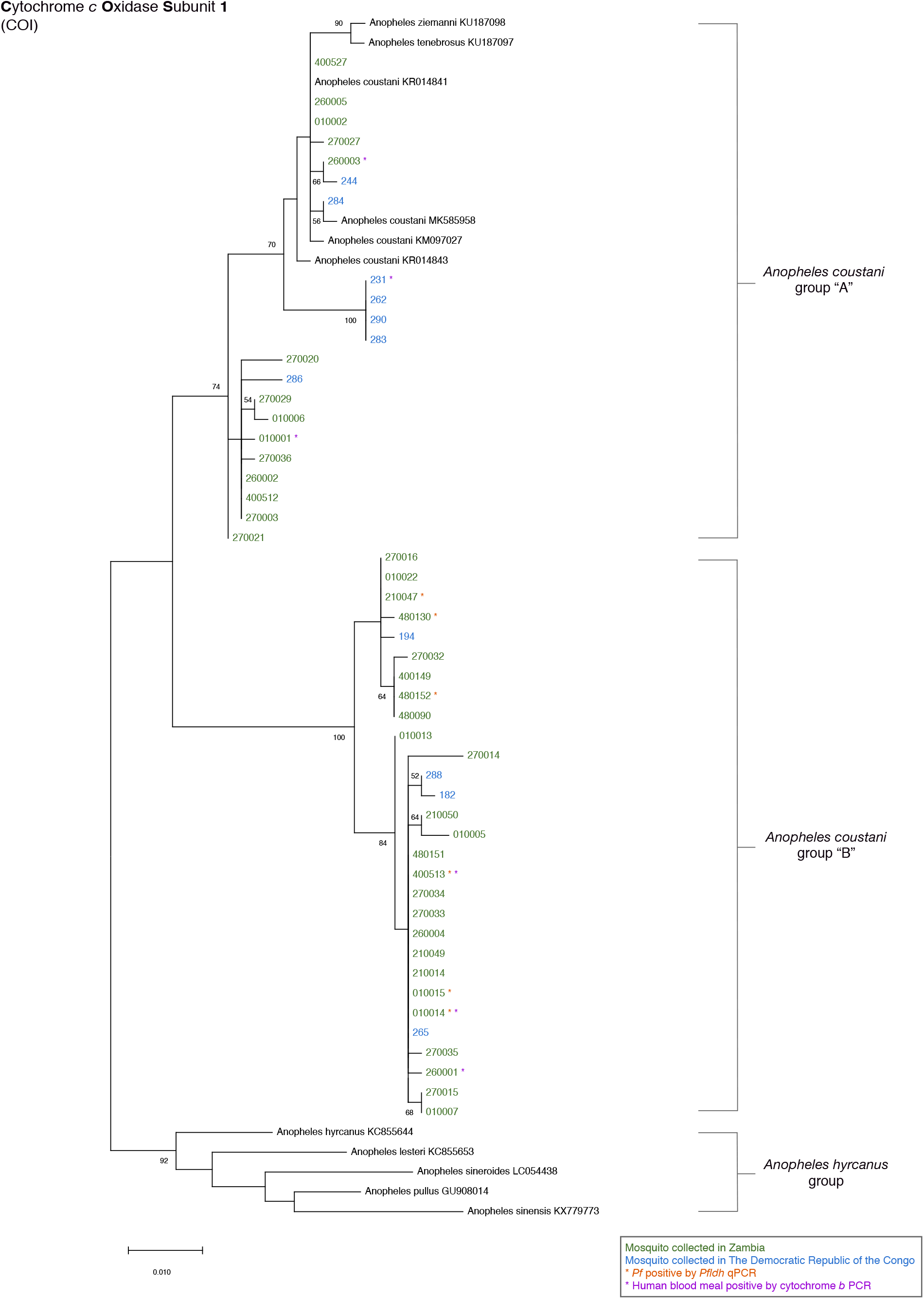
Cytochrome *c* oxidase subunit I (COI) Maximum Likelihood tree. This evolutionary analysis was performed using the Maximum Likelihood method and General Time Reversible model, with 1000 bootstraps for branch support (Nei and Kumar 2000). The tree with the highest log likelihood (−1603.26) is shown. The tree is drawn to scale, with branch lengths shown in the number of substitutions per site. This analysis contains 61 samples, including 50 samples from this collection, and 11 sample sequences obtained from NCBI BLASTn. Evolutionary analyses were conducted in MEGA X (Kumar et al. 2018, Stecher et al. 2020).

Similarly, the phylogenetic tree that was created for the ITS2 fragment (Figure 3) showed that all of the specimens morphologically identified as *An. coustani* clustered together. Moreover, the structuring revealed two groups (hereafter “*An. coustani* group A” and “*An. coustani* group B”) with 94% and 99% bootstrap support, respectively. *An. coustani* group A contained the same 21 sequences from field mosquitoes as *An. coustani* group A from the COI clustering, and the same was observed for the samples in the group B clusters for both genetic targets, providing a topological concordance between mtDNA COI and nuclear ITS2 phylogenetic trees. As observed for the COI target, previously published sequences of *An. coustani* clustered into *An. coustani* group A, even though they were from different geographical areas (KR014826 and KR014824 from Zambia, MK129245 from Madagascar, and KJ522815 from Kenya).

**Figure 3.**
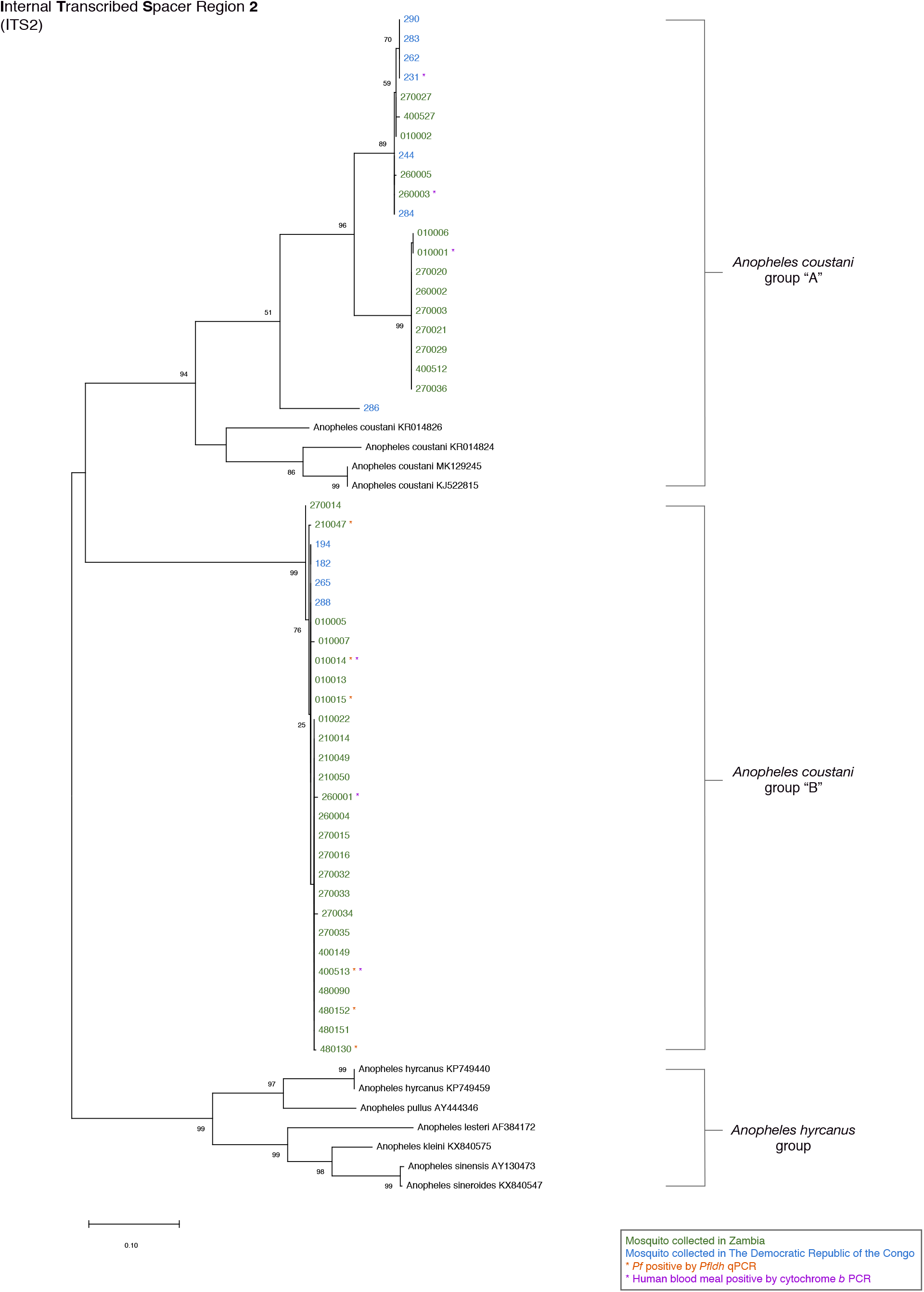
Ribosomal internal transcribed spacer region 2 (ITS2) Maximum Likelihood tree. This evolutionary analysis was performed using the Maximum Likelihood method and General Time Reversible model, with 1000 bootstraps for branch support (Nei and Kumar 2000). The tree with the highest log likelihood (−4723.20) is shown. The tree is drawn to scale, with branch lengths shown in the number of substitutions per site. This analysis contains 61 samples, including 50 samples from this collection, and 11 sample sequences obtained from NCBI BLASTn. Evolutionary analyses were conducted in MEGA X (Kumar et al. 2018, Stecher et al. 2020).

Genetic diversity parameters were calculated for each gene and *An. coustani* groups (Table 1). The mean haplotype diversity was higher for the *An. coustani* A groups of both targets (0.943±0.033 for COI and 0.762±0.050 for ITS2) when compared to the *An. coustani* B groups (0.860±0.038 for COI and 0.655±0.041 for ITS2). Nucleotide diversity was also higher for the *An. coustani* A groups according to both genetic targets (0.011 for COI and 0.052 for ITS2) when compared to the *An. coustani* B groups (0.007 for COI and 0.002 for ITS2) (Table 1). The low frequency of unique haplotypes in the *An. coustani* B groups is supported visually by both phylogenetic trees which generally show less clustering when compared to the A groups. Overall, haplotype and nucleotide diversity did not differ greatly between the two groups identified by the Maximum Likelihood phylogenies.

**Table 1.** Genetic diversity by genetic target and phylogenetic *An. coustani* group. DNAsp (v. 5.10.1) was used to calculate the number of polymorphic sites (S), the number of DNA haplotypes (h), the haplotype diversity (H_d_), the nucleotide diversity (π) among all loci, the average number of nucleotide differences (k), and theta per site from Eta.

|  | COI |  | ITS2 |  |
| --- | --- | --- | --- | --- |
|  | *An. coustani*<br>group A | *An. coustani*<br>group B | *An. coustani*<br>group A | *An. coustani*<br>group B |
| no. sequences | 21 | 29 | 21 | 29 |
| S (no. polymorphic sites) | 23 | 22 | 58 | 11 |
| h (no. haplotypes) | 14 | 17 | 7 | 7 |
| H<sub>d</sub> (haplotype diversity) ± Std. dev | 0.943±0.033 | 0.860±0.038 | 0.762±0.050 | 0.655±0.041 |
| π (nucleotide diversity) | 0.011 | 0.007 | 0.052 | 0.002 |
| k (average no. nucleotide differences) | 6.200 | 3.774 | 28.859 | 1.333 |
| θ (per site from Eta) | 0.012 | 0.009 | 0.024 | 0.004 |

## Discussion

All of the female *An. coustani* mosquitoes included in this study were caught outdoors, and some were found to be blooded either with human, goat, or a mixed blood meal of these two mammals. Although the sample size is small, these findings support reports that *An. coustani* is primarily exophagic (Nepomichene et al. 2015a, Degefa et al. 2017). While none of the female *An. coustani* mosquitoes in this study had a positive signal from CSP ELISA assays, six samples were qPCR-positive for *P. falciparum.* The qPCR is a more sensitive assay than ELISA, but the pitfall of this approach is that although it can specifically detect *P. falciparum* DNA, it cannot determine if it was from infectious sporozoites, early developmental stages, or even intact parasites. In this study, head and thoraces of the mosquitoes were used, which should restrict the presence of other parasite stages such as oocysts, but the limitation in this study is that the presence of gametocytes cannot be excluded (Carpi, unpublished). While this does not characterize the mosquitoes as infectious, finding human blood meals in 6 of 35 (3 solely human and 3 in combination with goat blood meal), and detection of *P. falciparum* DNA in 6 of 35 female *An. coustani* suggest that this mosquito feeds on human hosts frequently. Two of the samples that tested human-positive by PCR also tested qPCR-positive for *P. falciparum*.

Phylogenetic analyses of both COI and ITS2 fragments reveal that *An. coustani* from northern Zambia and southern DRC partition into two strongly supported groups, herein defined as *An. coustani* group A and *An. coustan*i group B. These groups have 100% bootstrap support in the COI analysis and 99% bootstrap support in the ITS2 analysis (Figures 2 and 3). As “*An. coustani”* likely comprises an undescribed species complex in which genetic structuring would not be unexpected, the biological significance of this observed structure is currently unknown. Importantly, the tree topology for both genetic targets which incorporates identical sample sets, strengthens the observed clustering of these specimens. Moreover, female mosquitoes in both *An. coustani* groups A and B (according to both targets) had taken human blood meals. It is of particular interest that two of the samples from *An. coustani* group B were found to have fed on human blood and/or goat blood and were also found to be qPCR-positive for *P. falciparum*, illustrating the convergence of human host, parasite and vector, although no causality can be proven here. Moreover, the presence of the identical haplotypes for both genetic targets at low frequency from northern Zambia and southern DRC (Figures 2 and 3) may suggest that inter-breeding and migrations might be occurring between the *An. coustani* subpopulations. However, larger sample sizes and further genetic investigation are required to confirm this scenario.

Given that *An. coustani* has been linked to malaria transmission in other regions of Africa and that specimens within this limited sample set are associated both with biting humans and acquiring *P. falciparum*, further studies in Zambia and the DRC are warranted (St. Laurent et al. 2016, Stevenson et al. 2016a). More extensive investigations are necessary to truly understand the foraging behavior and malaria transmission potential of *An. coustani* and other understudied vectors. Such vectors are easily overlooked and may be behaviorally resistant to the indoor-based vector interventions deployed across subSaharan Africa (Afrane et al. 2016). Behavioral resistance that may result from genetic adaptions or phenotypic plasticity in primary and alternative vectors (Govella et al. 2013). For instance, in the Solomon Islands, the scale-up of ITNs and IRS has successfully eliminated the primary vector *An. koliensis* and has left *An. punctulatus* with a fragmented distribution, allowing an exophagic species (*An. farauti*) to emerge as the new sole primary vector as it changed its behavior to avoid insecticides (Bugoro et al. 2011, Russell et al. 2013). These findings highlight the urgency to expand entomological surveillance to include other anopheline species, especially in transmission settings where primary endophagic and endophilic vector populations have been mostly successfully controlled by elimination strategies, but transmission still remains. In these regions, residual transmission will be maintained by mosquito populations that preferentially bite outdoors and are clear of indoor-based control measures. In these sub-Saharan settings, exophagic species such as *An. coustani* are ready to play a more significant role in transmission.

## Acknowledgements

The authors kindly acknowledge the Southern and Central Africa ICEMR field teams in Nchelenge District in Zambia and Kashobwe and Kilwa Health Zones in the DRC for their logistical assistance and participation in field collections. The authors are very grateful to the communities in Zambia and the DRC from which household collections were drawn and the National Malaria Control/Elimination Programs under the Ministries of Health in both DRC and Zambia.

## Author contributions

DEN and IIC conceived and developed the study. JCS, MM, GC, CMJ, and TK carried out field collections and compiled collection data. Samples were processed by IIC, CMJ, and TK and phylogenetic trees constructed and analyzed by IIC, GC, DEN and CMJ. IIC and DEN drafted the manuscript. All authors read and reviewed the final manuscript.

## References

Afrane, Y. A., M. Bonizzoni, and G. Yan. 2016. Secondary Malaria Vectors of Sub-Saharan Africa: Threat to Malaria Elimination on the Continent? In A. J. Rodriguez-Morales (ed.), Current Topics in Malaria. IntechOpen.

Altschul, S. F., T. L. Madden, A. A. Schäffer, J. Zhang, Z. Zhang, W. Miller, and D. J. Lipman. 1997. Gapped BLAST and PSI-BLAST: a new generation of protein database search programs. Nucleic Acids Res 25: 3389–3402.

Antonio-Nkondjio, C., Kerah CH, Simard F, Awono-Ambene P, Chouaibou M, Tchuinkam T, and F. D. 2006. Complexity of the malaria vectorial system in Cameroon: contribution of secondary vectors to malaria transmission. Journal of Medical Entomology 43: 1215–1221.

Beebe, N. W. 2018. DNA barcoding mosquitoes: advice for potential prospectors. Parasitology 145: 622–633.

Beier, J. 2002. Vector incrimination and entomological inoculation rates, pp. 3–11. In D. D. L. (ed.), Malaria methods and protocols, vol. 72. Humana Press, Methods in molecular medicine.

Bhatt, S., D. J. Weiss, E. Cameron, D. Bisanzio, B. Mappin, U. Dalrymple, K. E. Battle, C. L. Moyes, A. Henry, P. A. Eckhoff, E. A. Wenger, O. Briët, M. A. Penny, T. A. Smith, A. Bennett, J. Yukich, T. P. Eisele, J. T. Griffin, C. A. Fergus, M. Lynch, F. Lindgren, J. M. Cohen, C. L. J. Murray, D. L. Smith, S. I. Hay, R. E. Cibulskis, and P. W. Gething. 2015. The effect of malaria control on Plasmodium falciparum in Africa between 2000 and 2015. Nature 526: 207–211.

Bobanga, T., S. E. Umesumbu, A. S. Mandoko, C. N. Nsibu, E. B. Dotson, R. F. Beach, and S. R. Irish. 2016. Presence of species within the Anopheles gambiae complex in the Democratic Republic of Congo. Transactions of The Royal Society of Tropical Medicine and Hygiene 110: 373–375.

Bugoro, H., C. Iro’ofa, D. O. Mackenzie, A. Apairamo, W. Hevalao, S. Corcoran, A. Bobogare, N. W. Beebe, T. L. Russell, C.-C. Chen, and R. D. Cooper. 2011. Changes in vector species composition and current vector biology and behaviour will favour malaria elimination in Santa Isabel Province, Solomon Islands. Malaria Journal 10: 287.

Burkot, T. R., J. L. Williams, and I. Schneider. 1984. Identification of Plasmodium Falciparum-Infected Mosquitoes by a Double Antibody Enzyme-Linked Immunosorbent Assay*. The American Journal of Tropical Medicine and Hygiene 33: 783–788.

Chanda, E., M. Kamuliwo, R. W. Steketee, M. B. Macdonald, O. Babaniyi, and V. M. Mukonka. 2013. An Overview of the Malaria Control Programme in Zambia. ISRN Preventive Medicine 2013: 8.

Chizema-Kawesha, E., J. M. Miller, R. W. Steketee, V. M. Mukonka, C. Mukuka, A. D. Mohamed, S. K. Miti, and C. C. Campbell. 2010. Scaling up malaria control in Zambia: progress and impact 2005-2008. The American journal of tropical medicine and hygiene 83: 480–488.

Coleman, A. W. 2007. Pan-eukaryote ITS2 homologies revealed by RNA secondary structure. Nucleic Acids Res 35: 3322–3329.

Das, S., M. Muleba, J. C. Stevenson, D. E. Norris, and T. for the Southern Africa International Centers of Excellence for Malaria Research. 2016. Habitat Partitioning of Malaria Vectors in Nchelenge District, Zambia. The American Journal of Tropical Medicine and Hygiene 94: 1234–1244.

Degefa, T., D. Yewhalaw, G. Zhou, M.-C. Lee, H. Atieli, A. K. Githeko, and G. Yan. 2017. Indoor and outdoor malaria vector surveillance in western Kenya: implications for better understanding of residual transmission. Malaria journal 16: 443–443.

Degefa, T., A. Zeynudin, A. Godesso, Y. H. Michael, K. Eba, E. Zemene, D. Emana, B. Birlie, K. Tushune, and D. Yewhalaw. 2015. Malaria incidence and assessment of entomological indices among resettled communities in Ethiopia: a longitudinal study. Malaria Journal 14: 24.

Fang, Y., W.-Q. Shi, and Y. Zhang. 2017. Molecular phylogeny of Anopheles hyrcanus group members based on ITS2 rDNA. Parasites & Vectors 10: 417.

Ferrari, G., H. M. Ntuku, S. Schmidlin, E. Diboulo, A. K. Tshefu, and C. Lengeler. 2016. A malaria risk map of Kinshasa, Democratic Republic of Congo. Malaria journal 15: 27–27.

Fornadel, C. M., L. C. Norris, V. Franco, and D. E. Norris. 2011. Unexpected Anthropophily in the Potential Secondary Malaria Vectors Anopheles coustani s.l. and Anopheles squamosus in Macha, Zambia. Vector Borne and Zoonotic Diseases 11: 1173–1179.

Gillies, M. T., and B. DeMeillon. 1968. The Anophelinae of Africa south of the Sahara (Ethiopian Zoogeographical Region). Publications of the South African Institute for Medical Research 54.

Gillies, M. T., and M. Coetzee. 1987. A Supplement to the Anophelinae of Africa South of the Sahara. Johannesburg: South African Institute for Medical Research.

Gomes, F. M., B. L. Hixson, M. D. W. Tyner, J. L. Ramirez, G. E. Canepa, T. L. Alves e Silva, A. Molina-Cruz, M. Keita, F. Kane, B. Traoré, N. Sogoba, and C. Barillas-Mury. 2017. Effect of naturally occurring Wolbachia in Anopheles gambiae s.l. mosquitoes from Mali on Plasmodium falciparum malaria transmission. Proceedings of the National Academy of Sciences 114: 12566–12571.

Goupeyou-Youmsi, J., T. Rakotondranaivo, N. Puchot, I. Peterson, R. Girod, I. Vigan-Womas, M. O. Ndiath, and C. Bourgouin. 2019. Differential contribution of Anopheles coustani and Anopheles arabiensis to the transmission of Plasmodium falciparum and Plasmodium vivax in two neighboring villages of Madagascar. bioRxiv: 787432.

Govella, N. J., P. P. Chaki, and G. F. Killeen. 2013. Entomological surveillance of behavioural resilience and resistance in residual malaria vector populations. Malaria Journal 12: 124.

Hast, M. A., J. C. Stevenson, M. Muleba, M. Chaponda, J.-B. Kabuya, M. Mulenga, J. Lessler, T. Shields, W. J. Moss, D. E. Norris, f. t. Southern, and C. A. I. C. o. E. i. M. Research. 2019. Risk Factors for Household Vector Abundance Using Indoor CDC Light Traps in a High Malaria Transmission Area of Northern Zambia. The American Journal of Tropical Medicine and Hygiene 101: 126–136.

Jones, C. M., Y. Lee, A. Kitchen, T. Collier, J. C. Pringle, M. Muleba, S. Irish, J. C. Stevenson, M. Coetzee, A. J. Cornel, D. E. Norris, and G. Carpi. 2018. Complete Anopheles funestus mitogenomes reveal an ancient history of mitochondrial lineages and their distribution in southern and central Africa. Scientific Reports 8: 9054.

Kamuliwo, M., E. Chanda, U. Haque, M. Mwanza-Ingwe, C. Sikaala, C. Katebe-Sakala, V. M. Mukonka, D. E. Norris, D. L. Smith, G. E. Glass, and W. J. Moss. 2013. The changing burden of malaria and association with vector control interventions in Zambia using district-level surveillance data, 2006–2011. Malaria Journal 12: 437.

Kent, R. J., and D. E. Norris. 2005. Identification of mammalian blood meals in mosquitoes by a multiplexed polymerase chain reaction targeting cytochrome B. The American journal of tropical medicine and hygiene 73: 336–342.

Koukouikila-Koussounda, F., and F. Ntoumi. 2016. Malaria epidemiological research in the Republic of Congo. Malaria Journal 15: 598.

Kumar, S., G. Stecher, M. Li, K. Christina, and K. Tamura. 2018. MEGA X: Molecular Evolutionary Genetics Analysis across computing platforms. Molecular Biology and Evolution 35: 1547–1549.

Librado, P., and J. Rozas. 2009. DnaSP v5: a software for comprehensive analysis of DNA polymorphism data. Bioinformatics 25: 1451–1452.

Lobo, N. F., B. S. Laurent, C. H. Sikaala, B. Hamainza, J. Chanda, D. Chinula, S. M. Krishnankutty, J. D. Mueller, N. A. Deason, Q. T. Hoang, H. L. Boldt, J. Thumloup, J. Stevenson, A. Seyoum, and F. H. Collins. 2015. Unexpected diversity of Anopheles species in Eastern Zambia: implications for evaluating vector behavior and interventions using molecular tools. Scientific Reports 5: 17952.

Margoliash, E. 1963. Primary Structure and Evolution of Cytochrome C. Proc Natl Acad Sci U S A 50: 672–679.

Masaninga, F., E. Chanda, P. Chanda-Kapata, B. Hamainza, H. T. Masendu, M. Kamuliwo, W. Kapelwa, J. Chimumbwa, J. Govere, M. Otten, I. S. Fall, and O. Babaniyi. 2013. Review of the malaria epidemiology and trends in Zambia. Asian Pacific journal of tropical biomedicine 3: 89–94.

Messina, J. P., S. M. Taylor, S. R. Meshnick, A. M. Linke, A. K. Tshefu, B. Atua, K. Mwandagalirwa, and M. Emch. 2011. Population, behavioural and environmental drivers of malaria prevalence in the Democratic Republic of Congo. Malaria Journal 10: 161–161.

Mharakurwa, S., P. E. Thuma, D. E. Norris, M. Mulenga, V. Chalwe, J. Chipeta, S. Munyati, S. Mutambu, P. R. Mason, and I. T. Southern Africa. 2012. Malaria epidemiology and control in Southern Africa. Acta tropica 121: 202–206.

Ministry of Health Zambia, M. 2015. Zambia National Malaria Indicator Survey. Lusaka, Zambia: Ministry of Health., http://www.makingmalariahistory.org/wp-content/uploads/2017/06/Zambia-MIS2015_Jan20-nosigs.pdf.

MIS. 2019. Zambia National Malaria Indicator Survey PATH; National Malaria Elimination Centre.

Mohanty, A., S. Swain, S. K. Kar, and R. K. Hazra. 2009. Analysis of the phylogenetic relationship of Anopheles species, subgenus Cellia (Diptera: Culicidae) and using it to define the relationship of morphologically similar species. Infection, Genetics and Evolution 9: 1204–1224.

Moss, W. J., D. E. Norris, S. Mharakurwa, A. Scott, M. Mulenga, P. R. Mason, J. Chipeta, and P. E. Thuma. 2012. Challenges and Prospects for Malaria Elimination in the Southern Africa Region. Acta Tropica 121: 207–211.

Mukonka, V. M., E. Chanda, U. Haque, M. Kamuliwo, G. Mushinge, J. Chileshe, K. A. Chibwe, D. E. Norris, M. Mulenga, M. Chaponda, M. Muleba, G. E. Glass, and W. J. Moss. 2014. High burden of malaria following scale-up of control interventions in Nchelenge District, Luapula Province, Zambia. Malaria Journal 13: 153.

Mwangangi, J. M., E. J. Muturi, S. M. Muriu, J. Nzovu, J. T. Midega, and C. Mbogo. 2013. The role of Anopheles arabiensis and Anopheles coustani in indoor and outdoor malaria transmission in Taveta District, Kenya. Parasites & Vectors 6: 114–114.

Nambozi, M., P. Malunga, M. Mulenga, J.-P. Van Geertruyden, and U. D’Alessandro. 2014. Defining the malaria burden in Nchelenge District, northern Zambia using the World Health Organization malaria indicators survey. Malaria journal 13: 220–220.

Nardini, L., R. H. Hunt, Y. L. Dahan-Moss, N. Christie, R. N. Christian, M. Coetzee, and L. L. Koekemoer. 2017. Malaria vectors in the Democratic Republic of the Congo: the mechanisms that confer insecticide resistance in Anopheles gambiae and Anopheles funestus. Malaria Journal 16: 448.

Nei, M. 1987. Molecular evolutionary genetics, pp. 512. New York: Columbia University Press.

Nei, M., and S. Kumar. 2000. Molecular Evolution and Phylogenetics, Oxford University Press, New York.

Nepomichene, T. N. J. J., E. Tata, and S. Boyer. 2015a. Malaria case in Madagascar, probable implication of a new vector, Anopheles coustani. Malaria journal 14: 475–475.

Nepomichene, T. N. J. J., E. Tata, and S. Boyer. 2015b. Malaria case in Madagascar, probable implication of a new vector, Anopheles coustani. Malaria Journal 14: 475.

Norris, D. E., A. Shurtleff, Y. T. Touré, and G. C. Lanzaro. 2001. Microsatellite DNA polymorphism and heterozygosity among field and laboratory populations of Anopheles gambiae ss (Diptera: Culicidae). Journal of Medical Entomology 38(2): 336–340.

Norris, L. C., and D. E. Norris. 2015. Phylogeny of anopheline (Diptera: Culicidae) species in southern Africa, based on nuclear and mitochondrial genes. Journal of vector ecology: journal of the Society for Vector Ecology 40: 16–27.

Oaks SC Jr., M. V., Pearson GW, et al., editors. 1991. Vector Biology, Ecology, and Control. In I. o. M. U. C. f. t. S. o. M. P. a. Control (ed.), “Malaria: Obstacles and Opportunities”, vol. 7. National Academies Press (US).

Parr, J. B., C. Belson, J. C. Patel, I. F. Hoffman, P. Kamthunzi, F. Martinson, G. Tegha, I. Thengolose, C. Drakeley, S. R. Meshnick, V. Escamillia, M. Emch, and J. J. Juliano. 2016. Estimation of Plasmodium falciparum Transmission Intensity in Lilongwe, Malawi, by Microscopy, Rapid Diagnostic Testing, and Nucleic Acid Detection. The American Journal of Tropical Medicine and Hygiene 95: 373–377.

Pinchoff, J., H. Hamapumbu, T. Kobayashi, L. Simubali, J. C. Stevenson, D. E. Norris, E. Colantuoni, P. E. Thuma, and J. M. f. t. S. A. I. C. o. E. f. M. R. William. 2015. Factors Associated with Sustained Use of Long-Lasting Insecticide-Treated Nets Following a Reduction in Malaria Transmission in Southern Zambia. The American Journal of Tropical Medicine and Hygiene 93: 954–960.

PMI. 2019a. Zambia Malaria Operational Plan FY 2019. USAID: U.S. President’s Malaria Initiative.

PMI. 2019b. FY 2019 Democratic Republic of Congo Abbreviated Malaria Operational Plan. USAID: U.S. President’s Malaria Initiative.

Post, R. J., P. K. Flook, and A. L. Millest. 1993. Methods for the preservation of insects for DNA studies. Biochemical Systematics and Ecology 21: 85–92.

Russell, T. L., N. W. Beebe, R. D. Cooper, N. F. Lobo, and T. R. Burkot. 2013. Successful malaria elimination strategies require interventions that target changing vector behaviours. Malaria Journal 12: 56.

Sinka, M. E., M. J. Bangs, S. Manguin, Y. Rubio-Palis, T. Chareonviriyaphap, M. Coetzee, C. M. Mbogo, J. Hemingway, A. P. Patil, W. H. Temperley, P. W. Gething, C. W. Kabaria, T. R. Burkot, R. E. Harbach, and S. I. Hay. 2012. A global map of dominant malaria vectors. Parasites & vectors 5: 69–69.

St. Laurent, B., M. Cooke, S. M. Krishnankutty, P. Asih, J. D. Mueller, S. Kahindi, E. Ayoma, R. M. Oriango, J. Thumloup, C. Drakeley, J. Cox, F. H. Collins, N. F. Lobo, and J. C. Stevenson. 2016. Molecular Characterization Reveals Diverse and Unknown Malaria Vectors in the Western Kenyan Highlands. The American Journal of Tropical Medicine and Hygiene 94: 327–335.

Stecher, G., K. Tamura, and S. Kumar. 2020. Molecular Evolutionary Genetics Analysis (MEGA) for macOS. Molecular Biology and Evolution.

Stevenson, J., B. St. Laurent, N. F. Lobo, M. K. Cooke, S. C. Kahindi, R. M. Oriango, R. E. Harbach, J. Cox, and C. Drakeley. 2012. Novel Vectors of Malaria Parasite in the Western Highlands of Kenya. Emerging Infectious Diseases 18: 1547–1549.

Stevenson, J. C., L. Simubali, S. Mbambara, M. Musonda, S. Mweetwa, T. Mudenda, J. C. Pringle, C. M. Jones, and D. E. Norris. 2016a. Detection of Plasmodium falciparum Infection in Anopheles squamosus (Diptera: Culicidae) in an Area Targeted for Malaria Elimination, Southern Zambia. Journal of Medical Entomology 53: 1482–1487.

Stevenson, J. C., J. Pinchoff, M. Muleba, J. Lupiya, H. Chilusu, I. Mwelwa, D. Mbewe, L. Simubali, C. M. Jones, M. Chaponda, M. Coetzee, M. Mulenga, J. C. Pringle, T. Shields, F. C. Curriero, and D. E. Norris. 2016b. Spatio-temporal heterogeneity of malaria vectors in northern Zambia: implications for vector control. Parasites & Vectors 9: 510.

Stone, W., B. Grabias, K. Lanke, H. Zheng, E. Locke, D. Diallo, A. Birkett, M. Morin, T. Bousema, and S. Kumar. 2015. A comparison of Plasmodium falciparum circumsporozoite protein-based slot blot and ELISA immuno-assays for oocyst detection in mosquito homogenates. Malaria Journal 14: 451.

Sutcliffe, C. G., T. Kobayashi, H. Hamapumbu, T. Shields, S. Mharakurwa, P. E. Thuma, T. A. Louis, G. Glass, and W. J. Moss. 2012. Reduced Risk of Malaria Parasitemia Following Household Screening and Treatment: A Cross-Sectional and Longitudinal Cohort Study. PLOS ONE 7: e31396.

Wat’senga, F., E. Z. Manzambi, A. Lunkula, R. Mulumbu, T. Mampangulu, N. Lobo, A. Hendershot, C. Fornadel, D. Jacob, M. Niang, F. Ntoya, T. Muyembe, J. Likwela, S. R. Irish, and R. M. Oxborough. 2018. Nationwide insecticide resistance status and biting behaviour of malaria vector species in the Democratic Republic of Congo. Malaria journal 17: 129–129.

WHO. 2015. Global technical strategy for malaria 2016–2030. Geneva: World Health Organization.

WHO. 2019a. World malaria report 2019. Geneva: World Health Organization.

WHO. 2019b. High burden to high impact: a targeted malaria response. Geneva: World Health Organization.

